# A general approach for stabilizing nanobodies for intracellular expression

**DOI:** 10.1101/2021.04.06.438746

**Authors:** John Dingus, Jonathan C.Y. Tang, Constance Cepko

**Author notes:** currently at Columbia University, United States (Jonathan C.Y. Tang).

## Abstract

Conventional antibodies and their derived fragments are difficult to deploy against intracellular targets in live cells, due to their bulk and structural complexity. Nanobodies provide an alternative modality, with well documented examples of intracellular expression. Despite their promise as intracellular reagents, there has not been a systematic study of nanobody intracellular expression. Here, we examined intracellular expression of 75 nanobodies from the Protein Data Bank. Surprisingly, a majority of nanobodies were unstable in cells, illustrated by aggregation and clearance. Using comparative analysis and framework mutagenesis, we developed a general approach that stabilized a great majority of originally intracellularly unstable nanobodies, without compromising target-binding function. Additionally, we identified distinct sequence features impacting nanobody intracellular stability, and demonstrated mutationally stabilized nanobody expression in vivo, in the murine retina and in *E. coli*. This work will allow for standardized nanobody engineering for intracellular applications, potentiating a growing field of intracellular interrogation and intervention.

## Introduction

Nanobodies, amino terminal fragments derived from a special class of antibody lacking light chains^1^, are the smallest antibody derivatives that retain full antigen binding function. Composed of a single variable domain of the heavy chain (V_H_H), nanobodies boast several features that make them attractive tools for a range of applications. As monomers, they are versatile building blocks for protein engineering. Due to their compact binding interfaces, they have become invaluable as protein crystallization chaperones to resolve high resolution crystal structures^2,3,4^. Their modular serum half-life and superior tissue penetration are attractive characteristics for the development of therapeutic biologics^5,6^. Perhaps one feature that has been taken somewhat for granted is superior stability that facilitates nanobody intracellular expression. Full length antibodies, as well as bulkier antibody fragments, are not normally amenable to intracellular expression, partially owing to the reducing environment of the cytoplasm that prevents the formation of structurally crucial disulfide bonds. The ability to express nanobodies intracellularly opens the door for functional investigations of subcellular protein complexes and signaling pathways. Furthermore, intracellular nanobodies can be directed against novel therapeutic targets previously inaccessible to biologics.

Several groups have detailed intracellular nanobody expression to facilitate live imaging of subcellular factors^7,8,9^. However, few have commented on intracellular expression of nanobodies as a class as it pertains to stability. Those that have broadly detailed nanobody stability have done so in an extracellular context, in buffered solutions that do not resemble the cytoplasmic environment^13,14^. We have been developing tools for nanobody-mediated, fluorescence-based sorting of live, target expressing cells^10^, and genetic manipulation of cells expressing specific targets^11,12^. As we expanded our initial studies, we found that many nanobodies were not natively stable in the intracellular environment. We thus set out to investigate intracellular expression of nanobodies more systematically. To this end, 75 unique nanobody sequences from crystal structures uploaded to the Protein Data Bank (PDB) were cloned into mammalian expression vectors, fused C-terminally to a fluorescent protein (FP). Transfection and live-cell fluorescence imaging of these nanobody-FP fusions in both 293T and HeLa cells revealed that many nanobodies degrade and/or aggregate within the cytoplasm and nucleus, while others appear stable and soluble (Figure 1A). Following these observations, we set out to define intracellular instability based on sequence features, as well as to derive a standardized method for stabilization of previously intracellularly unstable nanobodies through framework mutagenesis. By leveraging positional sequence conservation apparent across intracellularly stable nanobodies, we have determined a method by which the vast majority of previously intracellularly unstable nanobodies can be stabilized for intracellular expression. This mutational stabilization, first observed in cell lines, was further validated for expression in vivo, in *E. coli*, and in the murine retina. These findings will contribute to the broader adoption of nanobodies as powerful reagents for both research and therapeutic applications.

**Figure 1.**
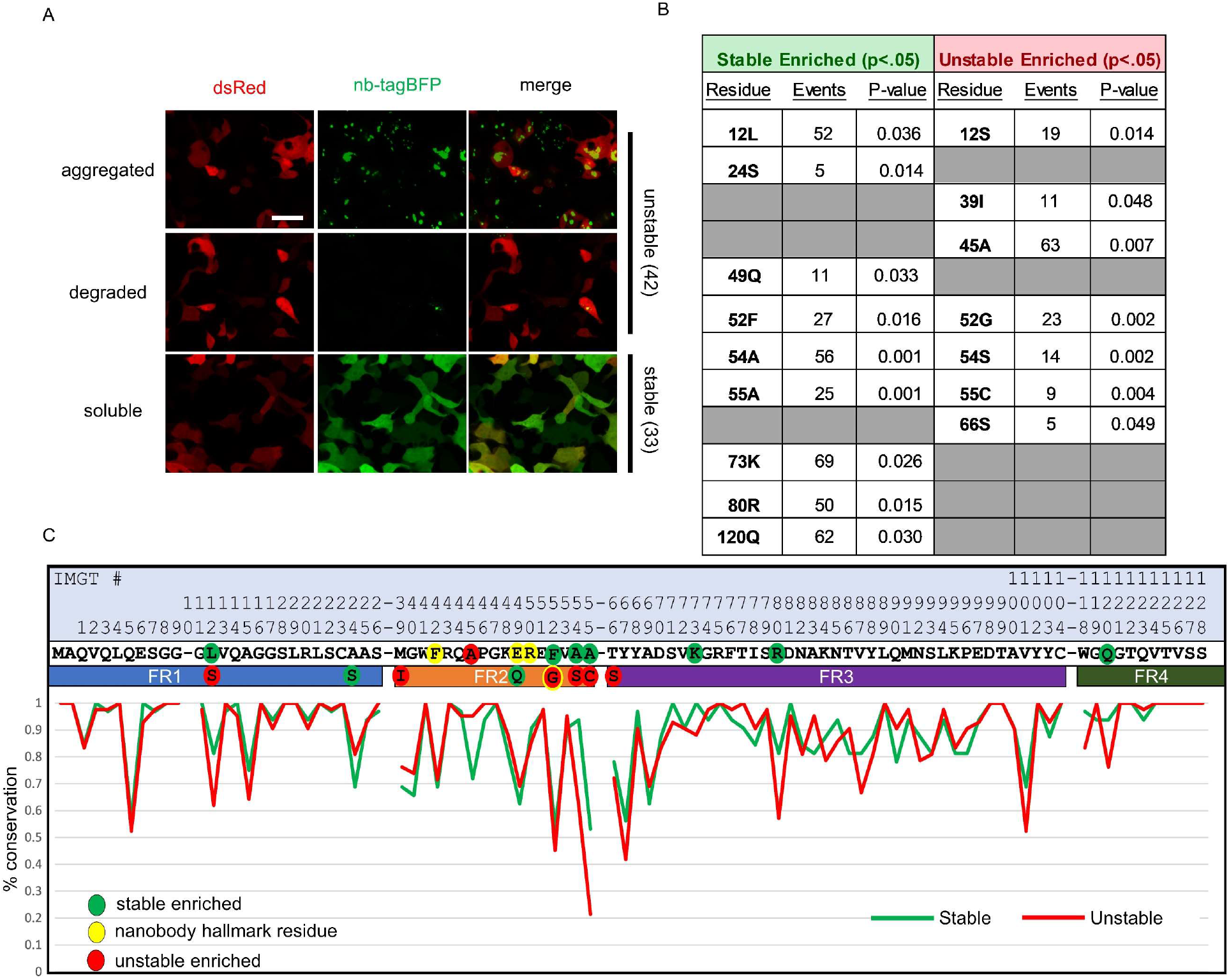
Sequence differences between intracellularly stable and unstable nanobodies. (A) Representative images of intracellularly stable and unstable nanobody-TagBFP fusions in transiently transfected 293T cells. Red signal is from co-transfected CAG-dsRed plasmid. (B) Significantly enriched positional residues between stable and unstable nanobodies (Fisher’s exact test). Events denote the total number of instances of each positional residue across all nanobodies (75 total). Grey cells denote positional insignificance. (C) Positional amino acid conservation across framework sequences of stable and unstable nanobodies. Scale bar is 20 μm.

## Results

### Classification of Intracellularly Stable and Unstable Nanobodies

To investigate nanobody intracellular expression broadly, we first compiled a repertoire of nanobody sequences to profile. The Protein Data Bank was combed for nanobody crystal structures and sequences, resulting in a list of 75 unique sequences, representing nanobodies derived from 3 camelid species and targeting 44 unique protein targets (sequences and targets detailed in Supplemental Table 1). These sequences were cloned into a mammalian expression vector in frame with TagBFP, linked by a 2 amino acid linker. All 75 sequences were expressed via transient transfection in both 293T and HeLa cells in separate wells of a 96 well plate. Intracellular expression patterns were captured by live fluorescence imaging. 33 out of 75 sequences exhibited strong, diffuse fluorescent signal, expected of well-expressed and intracellularly stable and soluble protein. Interestingly, 42 out of 75 sequences exhibited intracellular phenotypes suggestive of intracellular instability, including low or absent fluorescent signal coincident with varying degrees of aggregation (Figure 1A). These results were confirmed over several replicate experiments.

After binning the nanobodies into “stable” and “unstable” groups, their sequences were analyzed to identify features that distinguished the two groups. Nanobodies are composed of three variable loops (CDR1-3), responsible for the majority of target-specific interaction, and four framework regions (FR1-4), forming the conserved framework structure of the nanobody. First, the average length of variable loop CDR3 was calculated for each group, as CDR3 loop length is highly variable (between 5 and 26 amino acids for nanobodies in the examined repertoire) and CDR3 represents the area of both the greatest sequence-level and structural diversity across nanobodies. While there was little difference in average CDR3 loop length between the groups (16.4 amino acids for the stable group and 16.9 amino acids for the unstable group), one difference related to CDR3 stood out: 17/75 nanobodies contained a CDR3 cysteine that normally forms a disulfide bond tethering CDR3 to either CDR1 (first variable loop) or FR2 (second framework region); all 17 were in the unstable group (Table 1). Because disulfides do not form when nanobodies are expressed in the cytoplasm, this feature may represent a structural liability for intracellular expression.

**Table 1:**
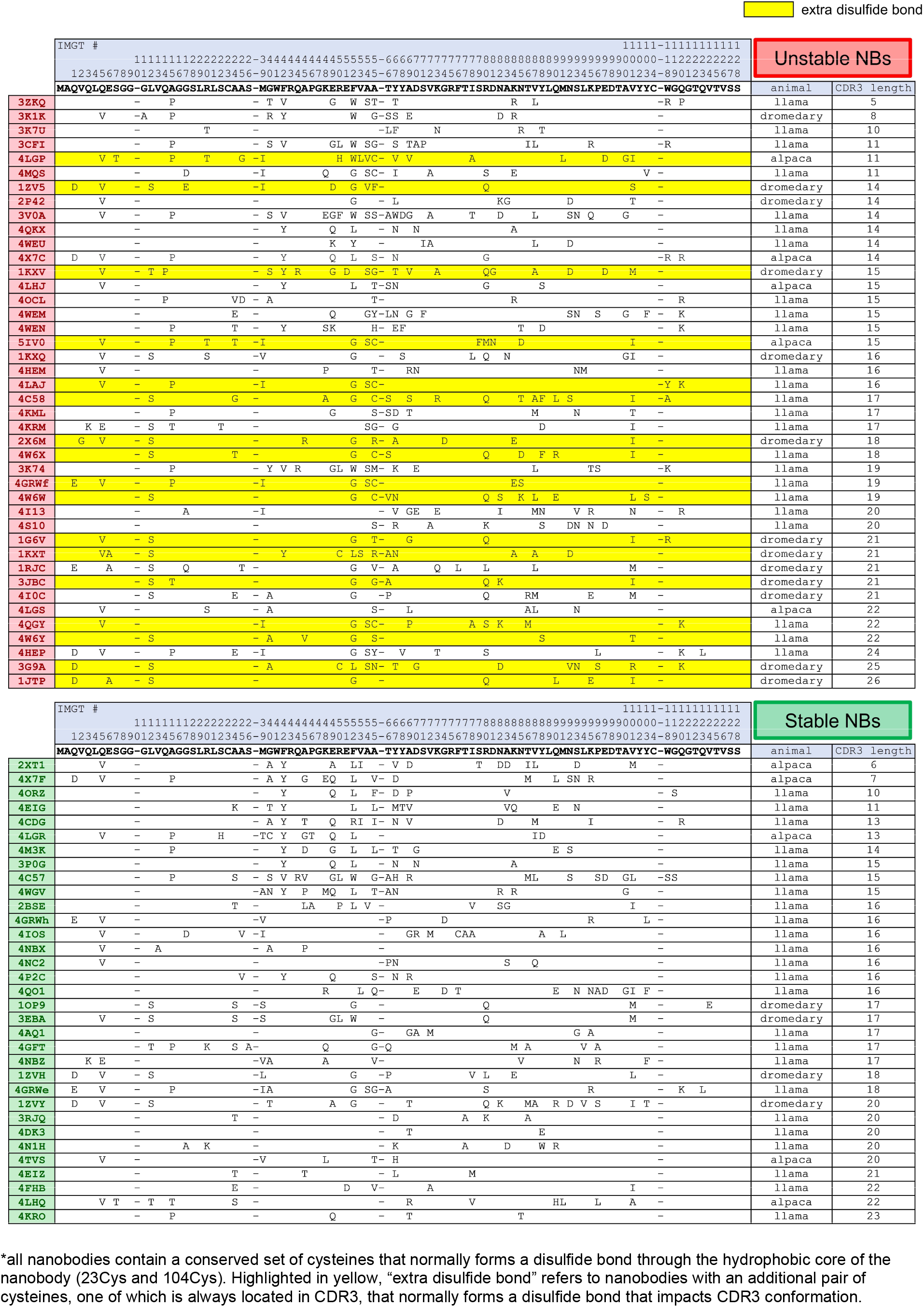
framework sequence variability (compared to consensus sequence) across stable and unstable nanobodies

We next derived the consensus sequence (the sequence constructed from the most common amino acid at each position across nanobody sequences) of the framework region (the region of sequence from which CDR sequences have been removed) for each group. Sequence level deviation between each stability-binned nanobody and the consensus sequence for stable nanobodies is illustrated in Table 1. Stable and unstable consensus sequences for the framework region were identical except at one position, amino acid 52 (IMGT numbering), where the most common amino acid across stable nanobodies was Phe and across unstable nanobodies was Gly. Interestingly, position 52 is one of four hallmark framework positions cited as differentiating VHs (heavy chain variable regions of conventional antibodies) and V_H_Hs^2^ (heavy chain variable regions of heavy-chain-only antibodies, analogous to nanobody). These four positions normally contribute to a hydrophobic interaction interface between VH and VL (light chain variable region) in conventional antibodies, and are substituted for amino acids that increase the hydrophilic character of the surface in V_H_Hs. 52Gly has been shown to enhance nanobody solubility, but at the expense of protein yields in *E. coli*, which may point to decreased stability^15,16^.

Next, we examined total positional enrichment for each amino acid across each position for both stable and unstable nanobody frameworks. Applying a Fisher’s exact test, 12 framework positions with residues that were significantly enriched in either group were identified, with 4 of those positions having a specifically enriched residue in both groups (Figure 1B). Of note, 52Gly was strongly enriched in the unstable group (19/23 occurrences in unstable nanobodies), as well as 55Cys, a cysteine that normally forms a disulfide bond with a CDR3 cysteine in a subset of nanobodies, as noted above (9/9 occurrences in unstable nanobodies). Additionally, 54Ser emerged as a strongly enriched unstable residue likely to drive instability (13/14 occurrences in unstable nanobodies). Position 54 points its functional group into the hydrophobic core of the nanobody, and is usually occupied by a small hydrophobic amino acid (93% of stable nanobodies examined contain 54Ala). A hydrophilic substitution at position 54 is likely to contribute to destabilization of the hydrophobic core.

Each unstable nanobody had at least 1 of 7 identified positional residues significantly enriched in the unstable group. However, the most common positional residue enriched in the unstable group, 45Ala, is the most common residue at that position for both groups, and is therefore unlikely to be a strong driver of instability (40/42 unstable nanobodies and 24/33 stable nanobodies contain 45Ala). 10/42 unstable nanobodies had 45Ala as their only enriched residue. Since this preliminary sequence analysis was unlikely to comprehensively explain the differences between stability groups, we decided to apply a broad mutagenesis strategy to unstable nanobodies to try to achieve stability.

### Stabilizing Mutagenesis Based on Positional Conservation

We set out to define a general mutagenesis approach to stabilize most, if not all, nanobodies for intracellular expression, taking advantage of sequence features observable across expression-profiled nanobodies. At the outset, our strategy was based upon the assumption that sets of important stabilizing residues are likely to be highly conserved across intracellularly stable nanobodies. While the framework consensus sequences for stable and unstable nanobodies are nearly identical, positional conservation varies between the two groups (Figure 1C). Our approach for conservation-based mutagenesis is schematized in Figure 2A. A threshold of ≥80% positional conservation was applied to generate a partial consensus sequence of the most highly conserved positional residues across stable nanobodies. This partial consensus framework was applied as a filter to identify non-conforming positional residues in each unstable nanobody for mutagenesis. At each position that a given unstable nanobody disagreed with the filter, that residue was changed to agree with the filter. An exception was made to maintain any cysteine that normally participates in a disulfide bond, as these cysteines likely impact positioning of CDR3. Applying these changes to all 42 intracellularly unstable nanobodies yielded mutants with a range of between 2 and 12 changes per nanobody, with an average mutational load of 5.7 changes per nanobody. Mutation numbers for each nanobody are reported in Tables 2B and C.

**Figure 2.**
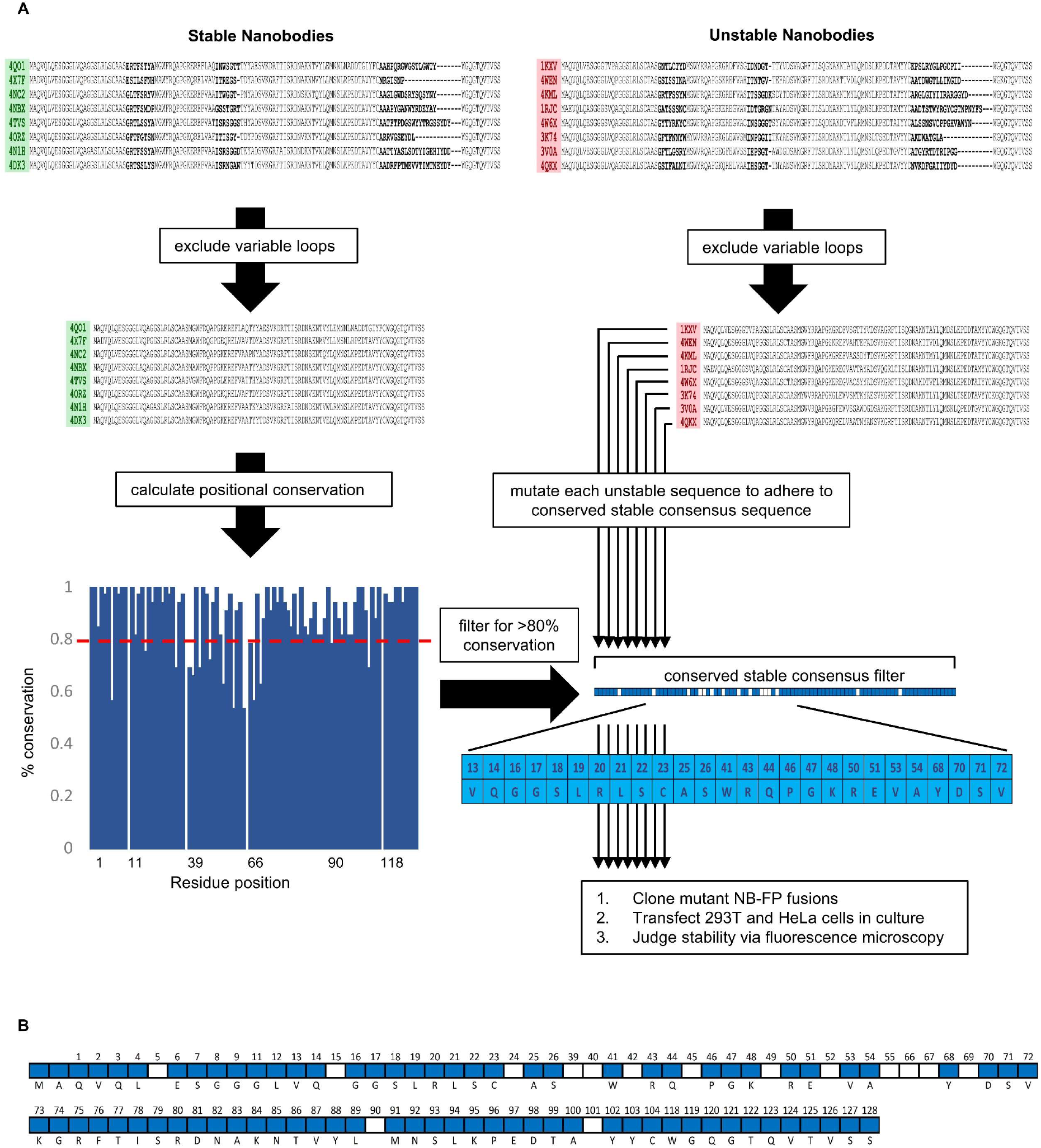
Schematic overview of conservation-based stabilizing mutagenesis strategy. (A) Nanobody-TagBFP fusions were classified as “stable” or “unstable” based on intracellular expression via transient transfection in 293T cells. Amino acid sequences were binned according to stability group, and variable domain sequences (CDRs) were excluded from downstream consideration. Positional sequence conservation was calculated across stable nanobody sequences, and positional amino acids of high conservation (>80%) were compiled to form a partial consensus filter. Each individual unstable nanobody framework was then compared to this filter, and any positional amino acid disagreement was resolved to adhere to the filter. Mutated nanobodies were then cloned, transfected into 293T and HeLa cells, and judged for stability via fluorescence microscopy. (B) All positional amino acids captured in the partial-consensus framework (blue cells). White cells denote framework positions excluded from the partial consensus framework.

**Table 2A:**
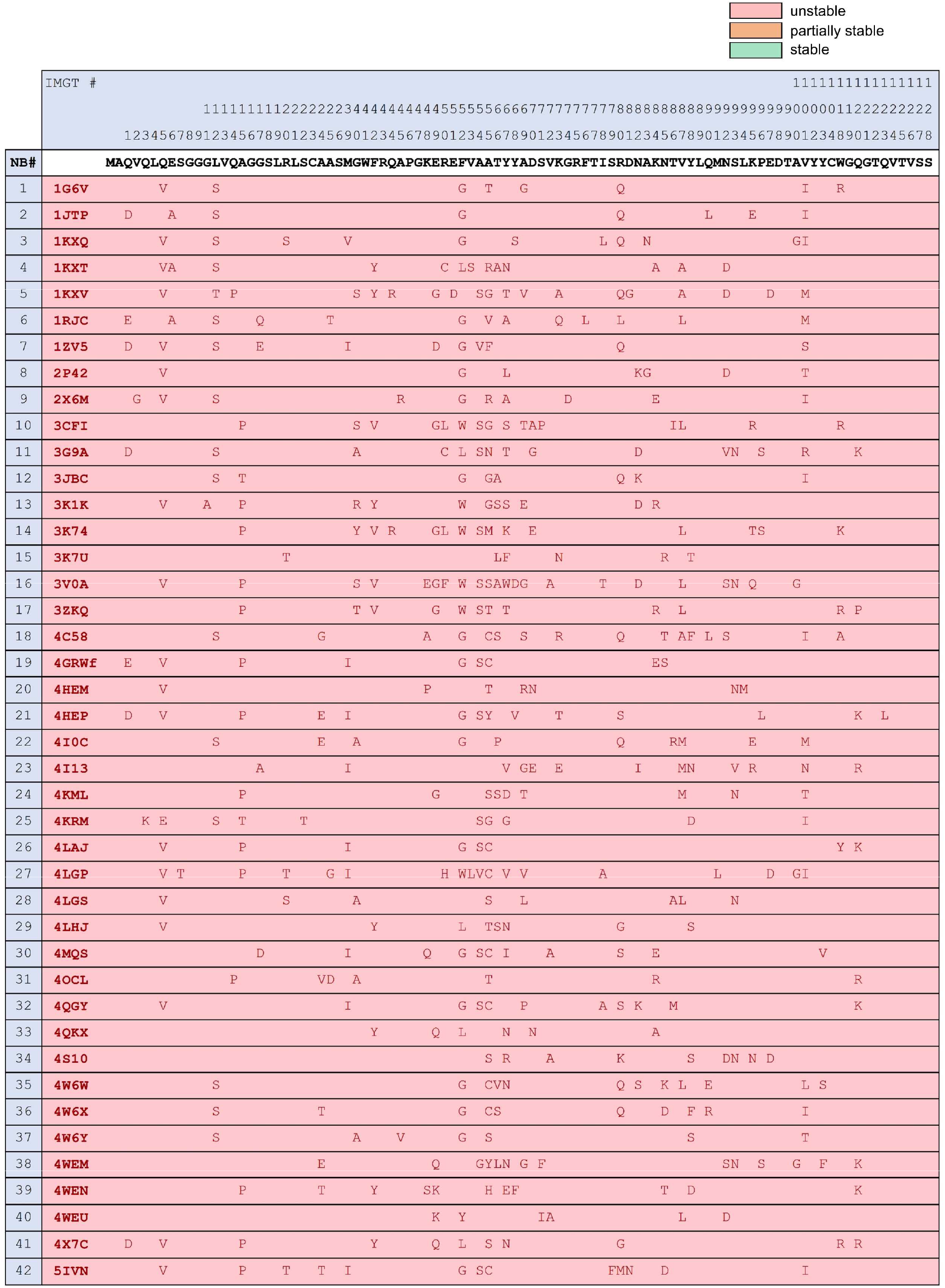
pre-mutagenesis framework sequence variability across unstable nanobodies

**Table 2B:**
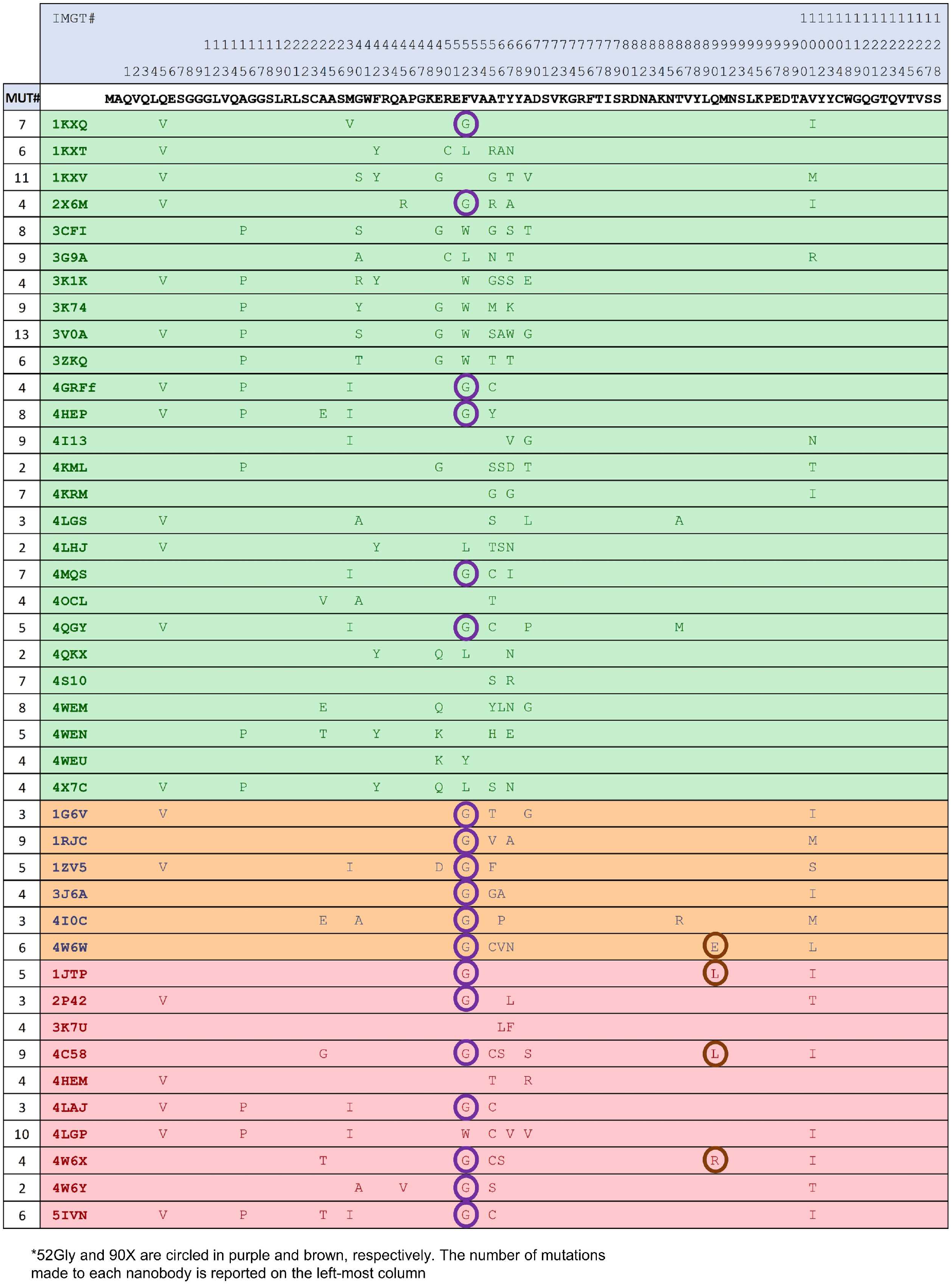
sequence and stability variability across nanobodies following stage 1 mutagenusis

**Table 2C:**
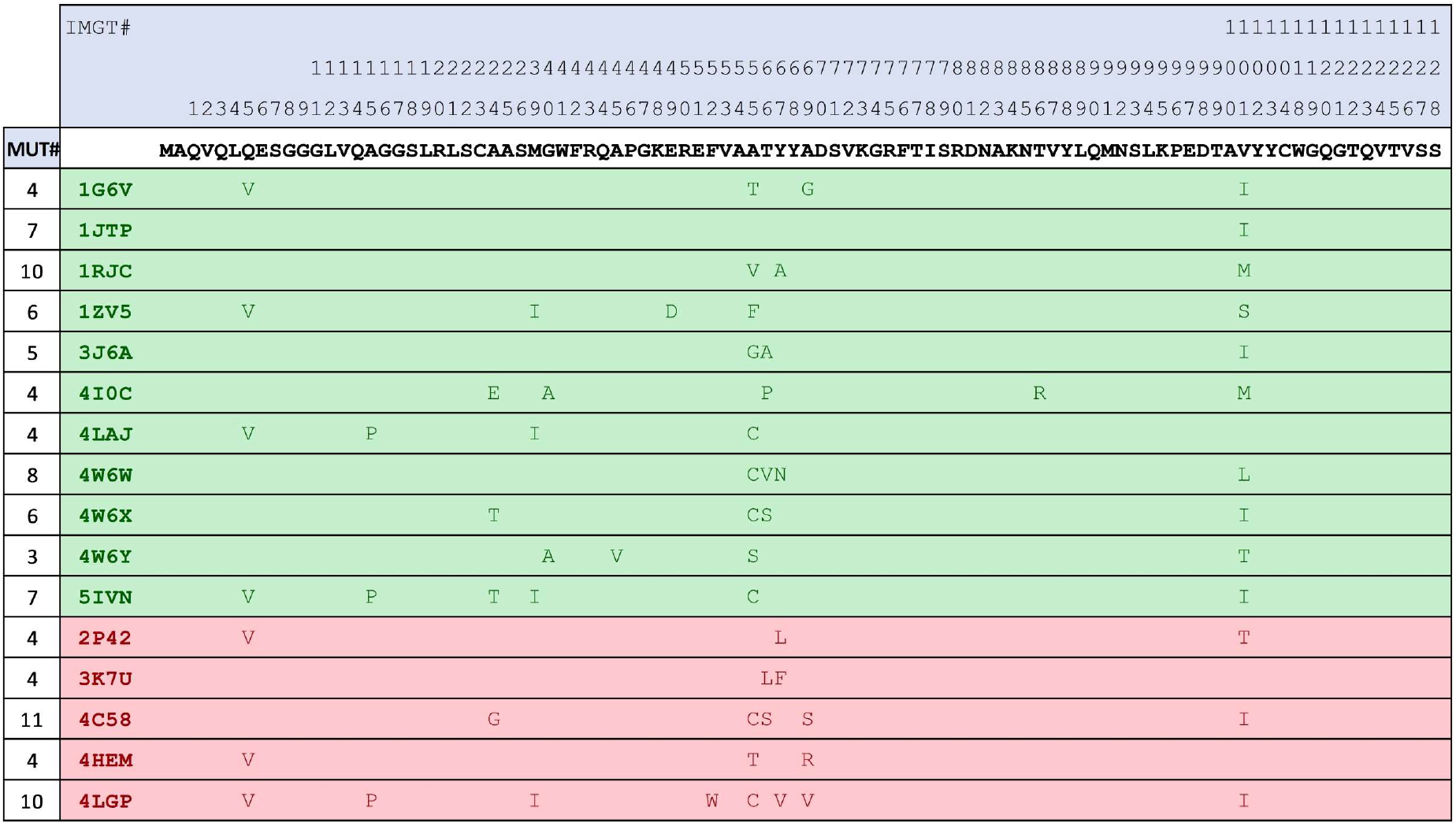
sequence and stability variability across nanobodies following stage 2 mutagenusis (+Gg2F + X90Q)

We cloned the derived mutant nanobodies and expressed them as FP fusions in 293T and HeLa cells as we had expressed their wild-type parents, previously. Sequence variability before and after mutagenesis for unstable nanobodies, as well as stability status following mutagenesis, is illustrated in Table 2. Fluorescent imaging revealed that 26/42 nanobodies had been effectively stabilized via targeted mutagenesis, as illustrated by strong fluorescent signal and a total absence of punctate aggregates (Figure 3A,C). Another 6/42 nanobodies exhibited improvement compared to their wild-type parent with respect to aggregation levels and soluble protein fraction. 10/42 nanobodies exhibited no improvement from mutagenesis.

**Figure 3.**
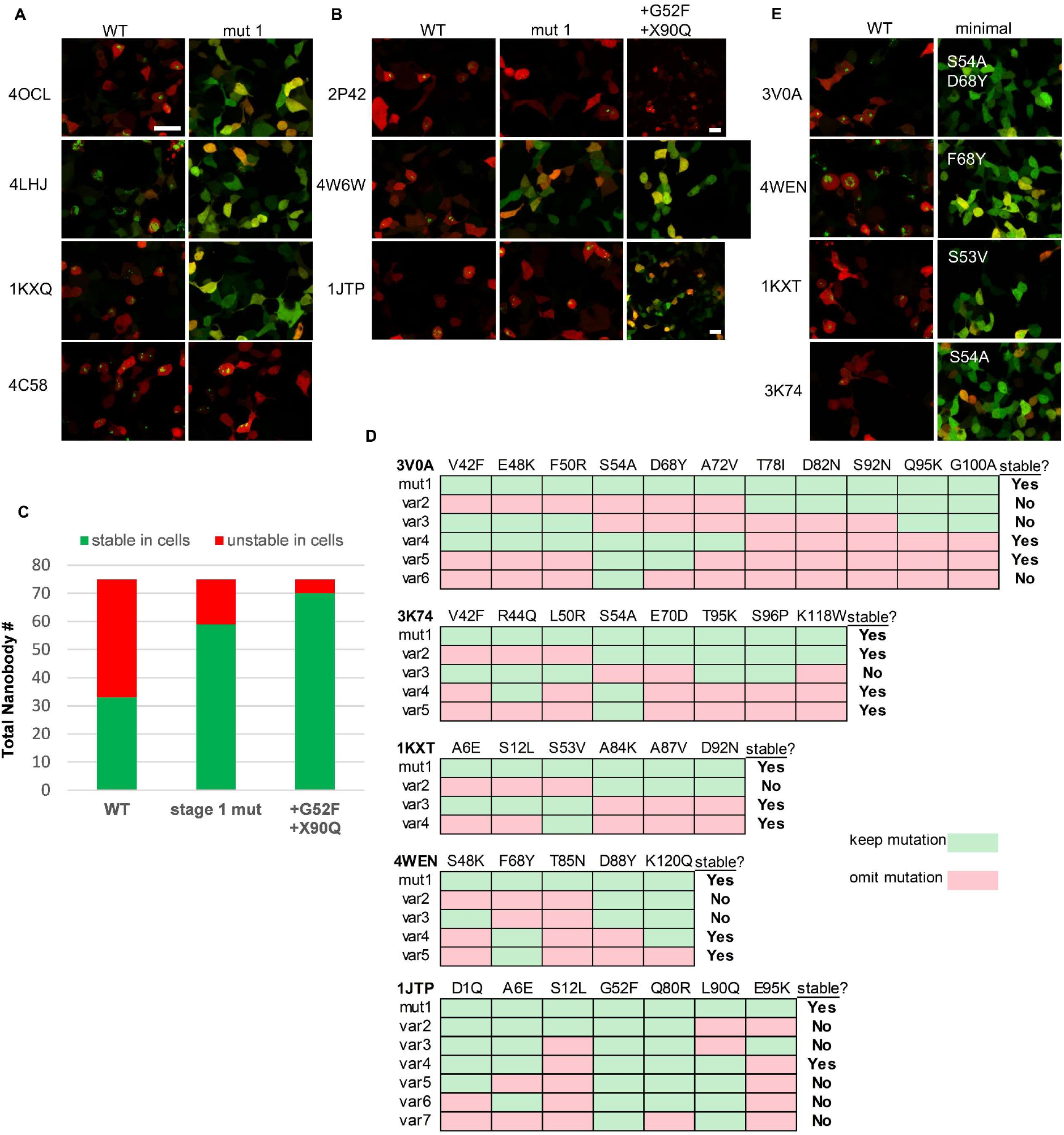
Mutationally stabilized nanobodies and specific stability drivers. (A) Examples of nanobody-TagBFP expression in 293T cells following transient transfection before (WT) and after (mut1) stage 1, conservation-based mutagenesis. Red signal is from co-transfected CAG-dsRed plasmid. (B) Examples of nanobody-TagBFP expression in 293T cells following transient transfection for nanobodies with Gly52. Wild-type, first-pass conservation-based mutants (mut1), and mutants with up to 2 additional mutations, Gly52Phe and X90Gln, are depicted. (C) Numbers of stable vs unstable nanobodies (75 total) following first-pass conservation-based mutagenesis (stage 1 mut) and final mutagenesis (+G52F +X90Q). (D) Example mutant variants tested for specific mutationally-stabilized nanobodies in order to identify necessary stability mutations. (E) Examples of nanobody-TagBFP expression in 293T cells following transient transfection for nanobodies effectively stabilized by 1-2 mutations. All scale bars are 20 μm.

Aligning the 42 mutant sequences, grouping sequences that were effectively, partially, or not stabilized via mutagenesis, a striking correlation emerged: 13/16 nanobodies that were not effectively stabilized via mutagenesis had 52Gly (Table 2B). Because position 52 is a relatively variable framework position, it was not included in the conservation filter for first pass mutagenesis. Additionally, non-consensus residues at positions 90 and 101 were enriched in non-stabilized nanobodies. For nanobodies not effectively stabilized by stage 1 mutagenesis, up to two additional changes were made wherever possible, Gly52Phe and X90Gln, to try to achieve stability. Because deviation at position 101 was almost perfectly correlated with the presence of Gly52, a likely driver of instability, this position was not changed (Table 2). As a result of this additional round of mutagenesis, 11/13 additional nanobodies were effectively stabilized. In total, 37/42 previously intracellularly unstable nanobodies were rendered stable and soluble (Figure 3C).

### Identification of Strong Drivers of Intracellular Stability

Encouraged by the success of the stabilizing mutagenesis approach, we set out to identify specific drivers of stability from amongst the partial consensus framework. To do this, we sought to eliminate “passenger” mutations for individual stabilized nanobodies and to identify the minimal, necessary mutational subsets required for intracellular stabilization. Five mutationally stabilized nanobodies with high mutational load and the high-confidence stability driver Ser54Ala were chosen for further examination (3K74, 3V0A, 3G9A, 4MQS, and 1KXV). 1KXT, a mutationally stabilized nanobody receiving the unique mutation Ser53Val, was also examined as both positions 53 and 54 point inward to the hydrophobic core of the nanobody. Subsets of the original mutations imposed to achieve stability were chosen, and new mutant variants were generated for intracellular stability testing. Mutation subsets were chosen based on crystal structure data and our own stabilization statistics. Several variants are depicted in Figure 3D and E. After testing several mutation variants for each nanobody, Ser54Ala was found to be sufficient to stabilize 3K74 on its own (originally stabilized with 8 mutations). Ser53Val similarly stabilized 1KXT by itself. 3V0A and 4MQS required only Ser54Ala and one additional mutation each to achieve stability (originally stabilized with 11 and 7 mutations, respectively). The minimal necessary sets of mutations needed to stabilize 3G9A and 1KXV were not identified, suggesting that several of their original mutations, in addition to Ser54Ala, were necessary for stabilization.

68Tyr and 80Arg were identified in nanobodies 3V0A and 4MQS (respectively) as important for their stabilization. These residues were subsequently investigated for their role in the stabilization of 3 additional nanobodies not originally containing the strong instability driver 54Ser: 4WEN, 4LHJ, and 1KXQ. 4LHJ was originally stabilized from only 2 mutations (Gln80Arg and Ser88Tyr). However, removal of Ser88Tyr did not destabilize 4LHJ, illustrating the stabilizing influence of 80Arg alone (data not shown). Surprisingly, 4WEN was effectively stabilized by 68Tyr alone, when it had originally received 5 mutations (Figure 3D and E). 1KXQ, which originally received both Ser68Tyr and Gln80Arg mutations in addition to 5 others, was effectively stabilized by 68Tyr alone, again illustrating the importance of 68Tyr for stability and suggesting only circumstantial importance of 80Arg (data not shown).

### Mutationally Stabilized Nanobodies Retain Target Binding Function in Cells

It is crucial for stabilizing mutagenesis to not only facilitate intracellular expression, but to maintain the nanobody’s ability to bind its target. Conventional antibodies engage their targets in a largely stereotyped fashion, relying heavily on variable loops for target interaction, with minimal direct framework contribution. In contrast, nanobodies exhibit a greater paratope diversity, and contribution of framework residues to binding is more common^17^. We compiled crystal structure data, curated by EMBL-EBI and made available through PDBe PISA, describing the interaction interfaces between nanobodies and their targets. Examination of these interfaces revealed that the great majority of interacting residues across nanobody framework regions are located in the most highly variable framework positions, positions omitted from our conservation mutagenesis approach (Figure 4A). While the possibility that mutation of non-interacting residues will generate conformational changes that impact target binding cannot be ruled out, this finding suggests that our approach likely minimizes impact on target binding, compared to strategies that rely on grafting variable loops onto established nanobody frameworks^18^.

**Figure 4.**
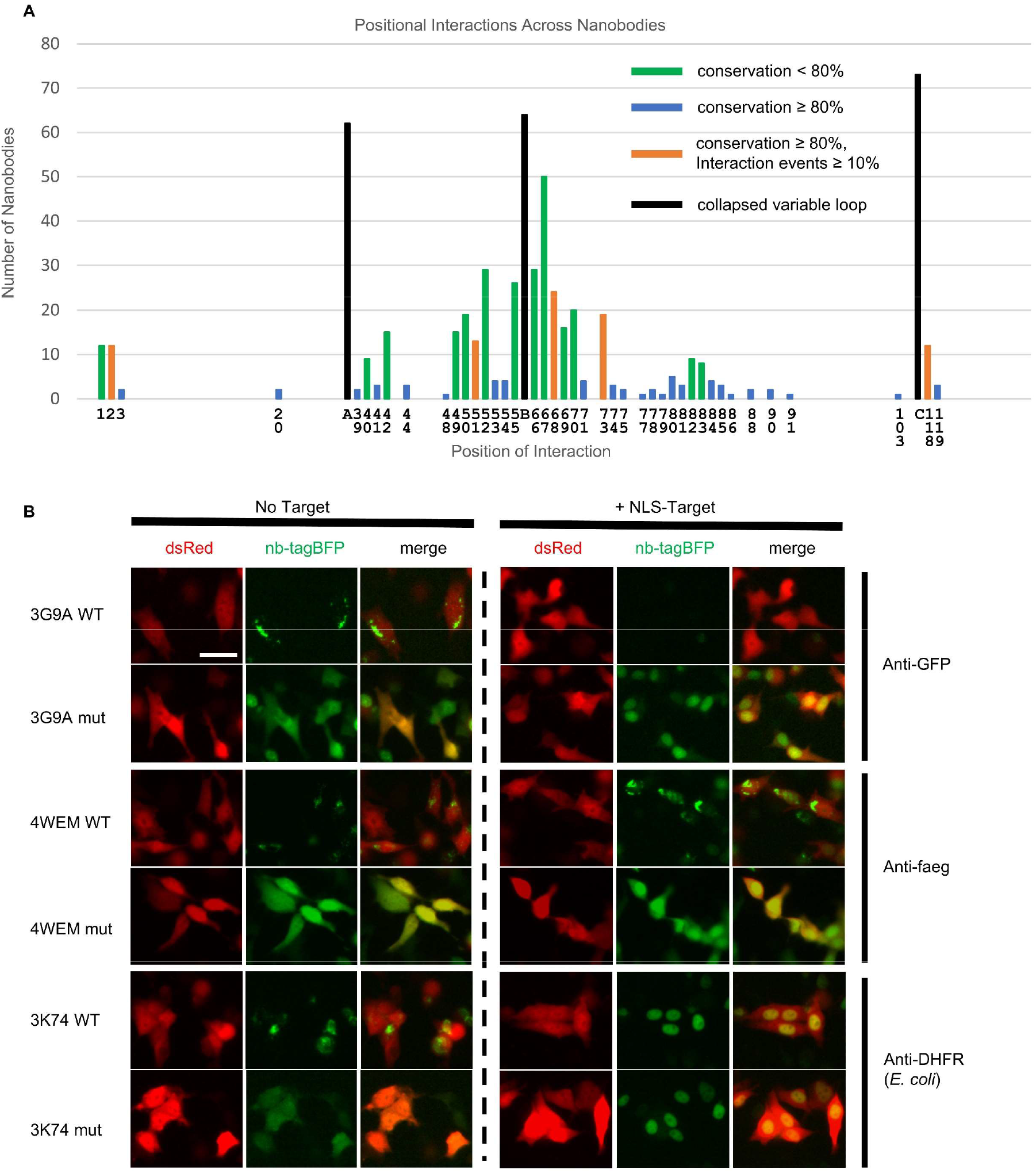
Target binding of parent and mutationally stabilized nanobodies. (A) Total number of target-interfacing positions across 75 nanobodies. Positions of interaction were determined based on buried surface area for crystal structure interfaces made available through PDBE-PISA. Black bars labelled “A, B, and C” denote positions across CDRs 1, 2 and 3, respectively. Colored bars describe sequence-level conservation across target-interfacing framework positions. (B) Representative images of nanobody-TagBFP expression in HeLa cells in the presence and absence of nuclearly localized-target. Wild-type and mutant nanobodies are depicted. Red signal is from co-transfected CAG-dsRed plasmid. Transfected DNA amount was normalized by addition of empty vector plasmid to transfection mix for the “no target” condition. Scale bar is 25 μm.

We chose 6 mutationally stabilized nanobodies whose targets are easily expressed intracellularly to test for intracellular target binding (3 targets, 2 nanobodies per target). Each nanobody was C-terminally linked with an FP, and each target was engineered to contain an N-terminal nuclear localization sequence (NLS). Wild-type and mutant nanobodies were transfected with either empty vector or NLS-target into 293T and HeLa cells. Target binding was assessed based on nuclear localization of nanobody-FP signal in the presence of NLS-target, observed via fluorescence imaging.

All 6 stabilized mutants were able to bind target (Figure 4B, Supplemental Figure 1). Of note was that 3G9A, which has a large CDR3 loop that is normally anchored by an extra disulfide bond to FR2, was able to bind target despite assumed loss of the disulfide bond. Interestingly, 2 nanobodies, 3K74 and 4I13, which are unstable in their wild-type forms in the absence of target, exhibited robust target binding in both their wild-type and mutationally stabilized forms. This suggested that the presence of their target, *E. coli* dihydrofolate reductase (DHFR), had a stabilizing effect (Figure 4B, Supplemental Figure 1). Our lab has previously described nanobody conditional stability in an engineered context^10^, and it is expected for a subset of nanobodies to exhibit this target-dependent stability naturally.

We next wanted to evaluate the necessity of using a partial consensus strategy for stabilization (Figure 2B), in which only highly conserved framework residues are fixed, as opposed to a full consensus strategy in which all framework residues are fixed (Figure 5A). The above 6 nanobodies were therefore tested for stability and target-binding after grafting their variable loops onto full consensus frameworks (50Cys was maintained for 3G9A). 6/6 nanobodies with full consensus frameworks exhibited stable expression in cells (Figure 5B). However, 2/6 nanobodies that achieved target binding in the context of our partial consensus framework (Figure 4B) lost binding in the context of the full consensus framework (Figure 5B). These nanobodies, 3K1K and 3K74, normally engage their targets along a concave framework interface (Figure 5C). Specifically, residues across FR2 form direct contacts with target, representing a common mode of binding (Figure 5C and D).

**Figure 5.**
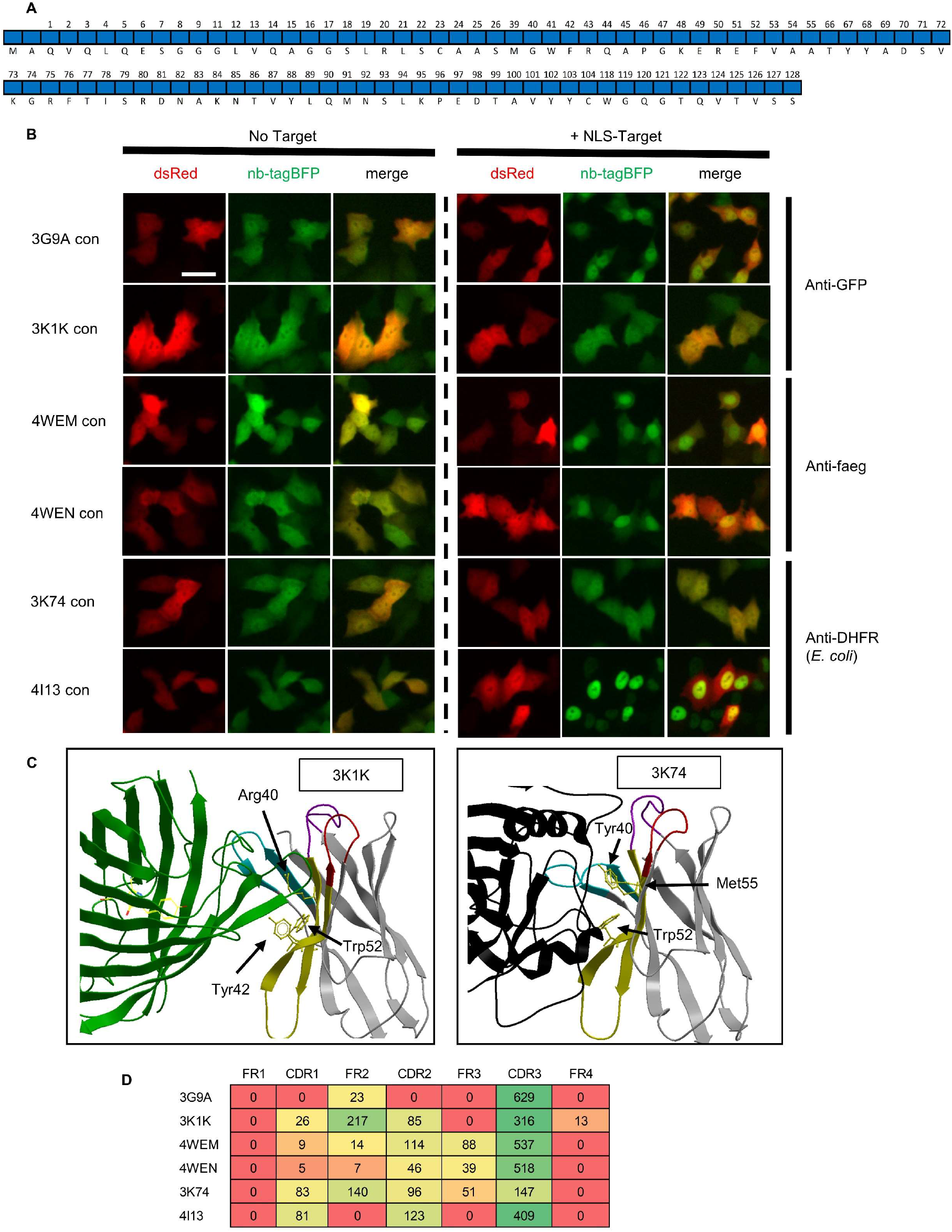
Target binding of nanobodies with full consensus frameworks. (A) All positional amino acids captured in the full consensus sequence. (B) Representative images of full consensus framework (con) nanobody-TagBFP expression in HeLa cells in the presence and absence of nuclearly localized-target. Red signal is from co-transfected CAG-dsRed plasmid. Transfected DNA amount was normalized by addition of empty vector plasmid to transfection mix for the “no target” condition. (C) Crystal structures of nanobodies that lose target binding when mutated to adhere to a full consensus framework are shown. Non-consensus framework residues directly contributing to target interface are depicted. (D) Target-interfacing surface area values in square angstroms (rounded to whole numbers) across distinct regions for nanobodies tested for target binding are shown. Values are taken from buried surface area interface values made available through PDBE-PISA. Scale bar is 25 μm.

We next turned our attention to a nanobody with an atypical mode of target binding. Nanobody 5IVN (also known as nanobody BC2) targets a short N-terminal stretch of β-catenin, representing a rare epitope: a short, linear peptide amenable for use as an affinity tag. Corroborating previous findings, we found 5IVN to be unstable when expressed in cells^7^. Unfortunately, while our mutagenesis approach stabilized 5IVN for cellular expression, it did not facilitate intracellular binding of its epitope as assessed by lack of co-localization with its target following cellular co-transfection of 5IVO-TagBFP and NLS-mCherry-epitope (data not shown). Binding was not achieved after limiting stabilizing mutagenesis to the single, necessary mutation to achieve intracellular stability, Ser54Ala. We suspect this lack of binding relates to loss of a crucial extra disulfide bond that normally orients its highly conformationally-precise paratope, but that does not form intracellularly. This disulfide bond has previously been shown to be required for target binding^19^.

Taken together, these target binding experiments suggest that nanobodies stabilized by the conservation-based mutagenesis approach developed here are likely to retain target binding function in cells, although certain conformational paratopes may not be amenable for intracellular recapitulation.

### Nanobody Intracellular Stability In Vivo

As many research and therapeutic applications of nanobody technology will require expression in vivo, we wanted to test whether improved nanobody intracellular stability established in cell lines would apply to in vivo settings. Electroporation of the retina in newborn mice is a means to deliver nanobody expression vectors to multiple cell types. Plasmids encoding wild-type or mutant 3V0A-TagBFP were injected into the subretinal space of postnatal day 2 (P2) mice, along with a dsRed control plasmid, and electroporation was carried out. Tissue was harvested at P12, and was fixed, sectioned, and imaged. Representative images are displayed in Figure 6A. Red fluorescent signal from the dsRed control plasmid delineated areas of successful electroporation. Little to no TagBFP signal was observed in cells electroporated with wild-type 3V0A-TagBFP. However, strong TagBFP signal was observed in cells electroporated with mutant 3V0A-TagBFP, mirroring observations in cultured cells. These cells included highly specialized sensory neurons (rod photoreceptors), interneurons (bipolar cells), and glia (Mueller glia), suggesting that mutationally stabilized nanobodies are stable in multiple, distinct cell types.

**Figure 6.**
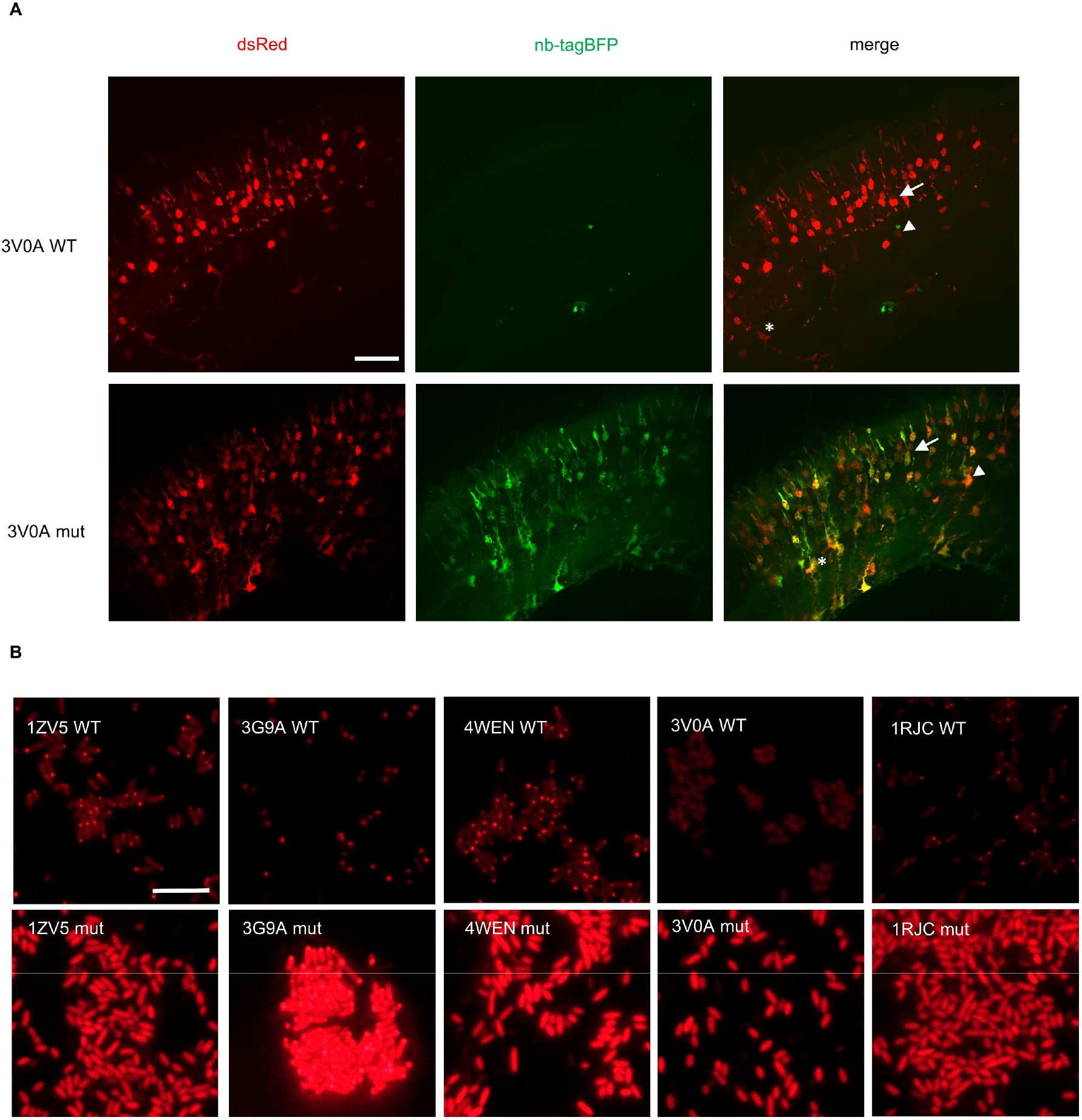
Nanobody Expression in the Murine Retina and *E. Coli*. (A) Representative Images of murine retina co-electroporated with CAG-dsRed and either wild-type or mutant CAG-3V0A-TagBFP plasmid. Retinas were electroporated on postnatal day 2 and harvested on postnatal day 12. Multiple cell types show expression following electroporation: photoreceptors (arrow), bipolar interneurons (triangle), and Mueller glia (asterix) are noted on merged images. (B) Representative images of BL21 *E. coli* transformed with either wild-type or mutant pRsetB-nanobody-mCherry plasmid. Images were taken following 4 hours of protein induction with 250 μM IPTG. All scale bars are 10 μm.

For additional context validation, we tested mutated nanobodies for stability in bacteria. Several groups have illustrated powerful applications taking advantage of nanobodies expressed in bacteria^20,21,22^. Bacteria offer a therapeutic modality, as they can be used as deliverable factories for nanobody production. Moreover, stable expression of nanobodies in bacteria would facilitate intra-bacterial studies, and may contribute to improved production yields for nanobody reagents produced in bacteria. Five mutationally stabilized nanobodies and their parents were tested for expression in BL21 *E. coli*. Nanobodies were fused C-terminally to mCherry and expression was assessed by fluorescence microscopy. All 5 wild-type nanobodies exhibited hallmarks of instability in bacteria (Figure 6B). 4/5 wild-type nanobodies exhibited protein aggregation as illustrated by focal inclusion bodies, in addition to diminished cytoplasmic fluorescence. 3V0A did not appear to aggregate, but exhibited low fluorescent signal. All 5 mutationally stabilized nanobody variants exhibited a marked increase in fluorescence level. 1ZV5, 4WEN, and 1RJC mutants showed no signs of protein aggregation, in contrast to their wild-type parents. Mutant 3G9A exhibited the most intense fluorescent signal, but also exhibited fluorescent inclusion bodies, possibly due to higher nanobody concentration.

## Discussion

Here, we describe a general and reliable method for modifying nanobody sequences to facilitate stable intracellular expression. A partial consensus framework was distilled from the most highly conserved positional residues across a large group of intracellularly stable nanobodies. This framework was applied broadly across unstable nanobodies to rescue intracellular stability in a great majority of cases. As new nanobodies are generated against intracellular targets, this stabilization approach should prove effective in achieving stable expression while maintaining target binding.

Consensus-based stabilization has been applied to antibody fragments in the past; Steipe and Wirtz applied consensus framework mutagenesis to V_L_ and V_H_ domains to enhance intracellular stability for conventional F_V_ fragments^23,24^. This approach was successful as conventional antibodies are amenable for loop grafting onto full donor frameworks due to their near exclusive reliance on CDRs for binding. Our partial consensus approach takes into account variable binding modes observed across nanobody crystal structures. Nanobodies often rely on direct framework engagement for target binding. Omitting framework positions of high variability from mutagenesis is important for preserving target binding in some, perhaps many, cases.

While investigating intracellular target binding, we noted that some nanobodies were able to effectively bind their targets despite their observed instability when target is absent. Such nanobodies may be useful reagents without the need of stability engineering in certain cases. However, in cases of high nanobody expression and low target availability, an unstable, aggregation-prone fraction of unbound nanobody may prove problematic for studies concerning subcellular protein localization, or for reasons of cellular toxicity. Indeed, one potential therapeutic application for intracellular nanobodies is to block protein aggregation that leads to various neurodegenerative diseases^25^. Several groups have investigated nanobodies as potential reagents for blocking aggregation of a range of cytotoxic, aggregation-prone factors including prion protein, amyloid beta, and alpha synuclein^26,27,28^. In these contexts, nanobody stability and solubility may prove critical to ensuring that no undue cellular stress is imparted by the treatment agent.

The potential for intracellular deployment of nanobody-based reagents is still far from being realized. Taking full advantage of the modularity and versatility inherent of these pared-down structures will require methods such as those proposed here to condition nanobodies for the intracellular environment.

## Materials and Methods

### Compilation of nanobody sequence and structure data

Nanobody sequences were pulled directly from the PDB database (sequences for this study compiled in 2016). To generate an interface atlas profiling positions of interaction between each nanobody and its target (linked data provided), interface data were referenced from the Proteins, Interfaces, Structures, and Assemblies tool (PDBePISA), provided by the European Bioinformatics Institute (EMBL-EBI). Values for positional interactions were taken from Buried Surface Area values (Å^2^) for each residue.

### Generation and cloning of nanobody sequences

Nanobody sequences were synthesized as double stranded DNA fragments (gBlocks) by IDT, and cloned directly into a CAG expression plasmid (Addgene plasmid 11150) (Matsuda and Cepko, 2004) in frame and linked with TagBFP via Gibson assembly (New England Biolabs, E2611). For bacterial expression constructs, nanobody sequences were PCR amplified with primers containing terminal regions of homology for cloning into a pRset plasmid (Addgene plasmid 3991)(Invitrogen, V35120) in frame and linked to mCherry. DH5α *E. coli* were transformed with assembled DNA and cultured in 4 mL cultures overnight in LB medium with 100 ug/mL carbenicillin. Plasmid DNA was purified using Qiagen miniprep kits.

Mutant nanobody sequences were designed by comparing intracellularly unstable nanobodies to a partial consensus framework sequence of stable nanobodies, and changing amino acids in each unstable nanobody to match the partial consensus sequence at each position. Nanobody framework regions were defined by The International Immunogenetics Information System (IMGT), a global reference for immunogenetics. The consensus sequence for stable nanobodies was generated by calculating the most frequent amino acid at each framework position across 33 intracellularly stable nanobodies. The partial consensus sequence represents the subset of consensus positions at which 80% or greater of the 33 stable nanobodies had the same amino acid (Figure 2B).

### Cell culture and transfection

All constructs used in mammalian cell transfection experiments were cloned into the pCAG plasmid. Plasmid DNAs encoding nanobody-TagBFP sequences were transfected into both 293T cells and HeLa cells, separately. Experiments were first conducted in 293T cells, validated across 3 replicates, and later verified in HeLa cells. 1 day prior to plasmid transfection, cells were plated in black, clear bottomed 96-well plates (CLS3603, Sigma-Aldrich) at roughly 10,000 cells/well and incubated at 37°C, and 5% CO_2_. For transfection of cells in each well, 5 uL of serum-free media was added to 200 ng of plasmid DNA (100 ng CAG-nanobody-TagBFP plasmid and 100 ng CAG-dsRed plasmid (Addgene plasmid 11151) (Matsuda and Cepko, 2004)). For target-binding experiments, an additional 100 ng of NLS-target expressing plasmid or 100 ng of empty vector (control) was added. 1 uL (or 1.5 uL for target-binding experiments) of 1 mg/mL PEI (Polysciences, 24765-2) dissolved in water was added to DNA in media and vortexed for 10 seconds. Resulting transfection mix was left to sit at room temperature for 10 minutes before being added to cells.

### Live fluorescent imaging and stability scoring

For initial stability scoring, 75 nanobodies were transfected into HEK293T cells (as described above) as TagBFP fusions driven by a CAG promoter. Roughly 24 hours after transfection, TagBFP signal was evaluated using a Leica DMI3000B microscope and a 20X objective lens. CAG-dsRed signal served as both a transfection control and orienting signal to assess cellular morphology. The 75 nanobodies were binned broadly into “stable” and “unstable” groupings based on the observed character of the TagBFP signal for each nanobody. Nanobodies with TagBFP signal that filled cells, mirroring dsRed signal, and that exhibited, at most, only minor and infrequent fluorescent puncta, were binned into the “stable” group. Nanobodies with sparse to absent TagBFP signal, and/or that exhibited major and frequent fluorescent puncta, were binned into the “unstable” group (Figure 1A). Groupings were validated across 3 separate rounds of transfection and evaluation in HEK293T cells, and further validated in HeLa cells. Grouping criteria were assigned to capture major differences in intracellular expression between groups, rather than subtle differences in expression pattern or character within groups.

For target-binding experiments, images were taken using the same Leica DMI3000B microscope and 20X objective lens roughly 24 hours after transfection. Excitation time was kept consistent between samples.

Imaging of parent and mutant nanobodies to evaluate mutational stabilization was carried out via automated confocal imaging. Using a PE Opera high-throughput confocal imaging system (https://www.flyrnai.org/supplement/BRO_OperaHighContentScreeningSystem.pdf) provided by the Drosophila RNAi Screening Center (https://fgr.hms.harvard.edu/), live, transfected HEK293T cells were imaged in 96-well plates using a water-immersion 40X objective. 10 fields for image acquisition, standardized across each well-plate well, were assigned prior to imaging. Through an automated protocol, 6 Z-stacks were taken in each field in both red and blue channels, spanning 12 microns. Following image acquisition, max projection images were generated, and nanobody-TagBFP signal was evaluated. in both red and blue channels, spanning 12 microns. Following image acquisition, max-projection images were generated, and nanobody-TagBFP signal was evaluated.

### Retinal electroporation

Wild-type and mutant CAG-3V0A-TagBFP plasmids were each electroporated along with control CAG-dsRed plasmid (1.5 ug/mL DNA) into the retina of P2 mouse pups. 3 pups received wild-type 3V0A plasmid, and 3 pups received mutant 3V0A plasmid. Electroporation was executed according to the methodology described in Matsuda and Cepko, 2004.

### Retinal histology and imaging

Electroporated retinas were harvested at P12. Retinas were fixed in 4% formaldehyde for 30 minutes, transferred to PBS for 10 minutes, and transferred to 15% sucrose in PBS for 30 minutes, all at room temperature. Fixed retinas were then embedded in OCT, flash frozen, and stored at −80°C. 12 μm sections were made using a cryostat, placed on glass slides, and mounted with Fluoromount-G mounting media (ThermoFisher, 00-4958-02). Images were taken using a Nikon Ti2 inverted microscope (spinning disk confocal) with a 40X oil immersion objective lens.

### Bacterial expression and imaging

Nanobody sequences were cloned into the bacterial expression vector pRsetB in frame and fused to mCherry. Plasmid was transformed into BL21(DE3) *E. coli* (New England Biolabs, C2527), and individual colonies were picked into 5 mL LB and cultured overnight at 37°C and shaken at 250 RPM. In the morning, 1 mL of each culture was added to 4 mL of M9 minimal media, and IPTG was added to a final concentration of 250 μM to induce protein expression. Induction cultures were incubated for 4 hours at 37°C and 250 RPM. 40 uL of induced culture was pipetted onto 3% M9-agar on a glass slide and covered with a glass coverslip. Bacteria were imaged with a Nikon Eclipse e1000 microscope using a 60X oil immersion objective lens. All images were taken with consistent excitation time. Experiments were validated across 3 replicates.

### Image processing

Images were processed using ImageJ. Images from Figures 1 and 3 received the following processing and no other adjustments: 1) images were converted to 8-bit, 2) Z stacks for each channel (red and blue) were merged to create maximum Z projections, 3) maximum Z projections for each channel were merged with one another, with red images inserted into the red channel, and blue images inserted into the green channel for greater contrast. Retinal images (Figure 6) received the same processing, but received additional contrast adjustment (all images received consistent adjustment to contrast). Other images, including bacterial images (images from Figures 4–6) without Z stacks were similarly converted to 8-bit, were color-channel merged, and received contrast equally administered contrast adjustment.

## Supplement

**Supplemental Table 1:**
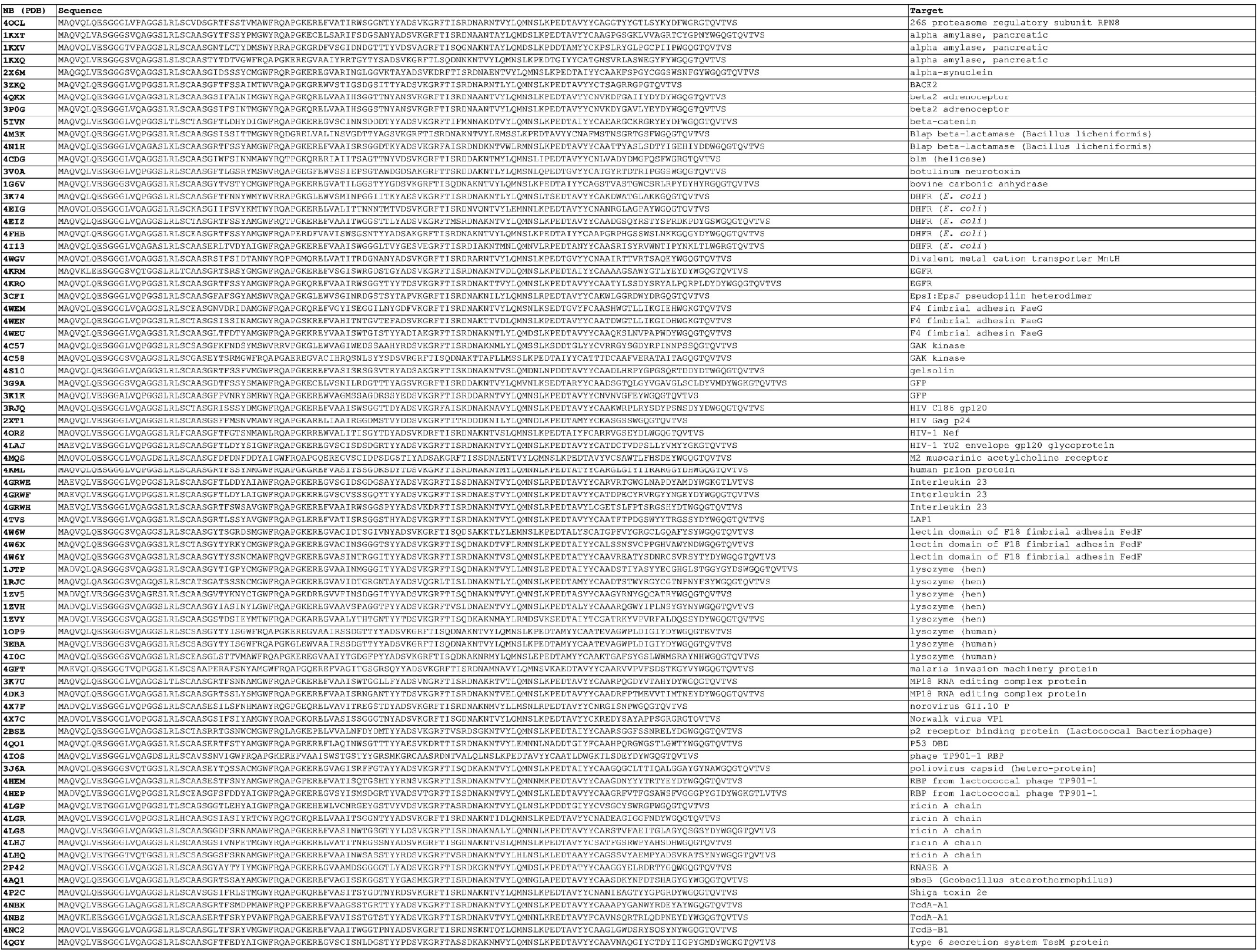
list of nanobodies and their associated antigens

**Supplemental Figure 1.**
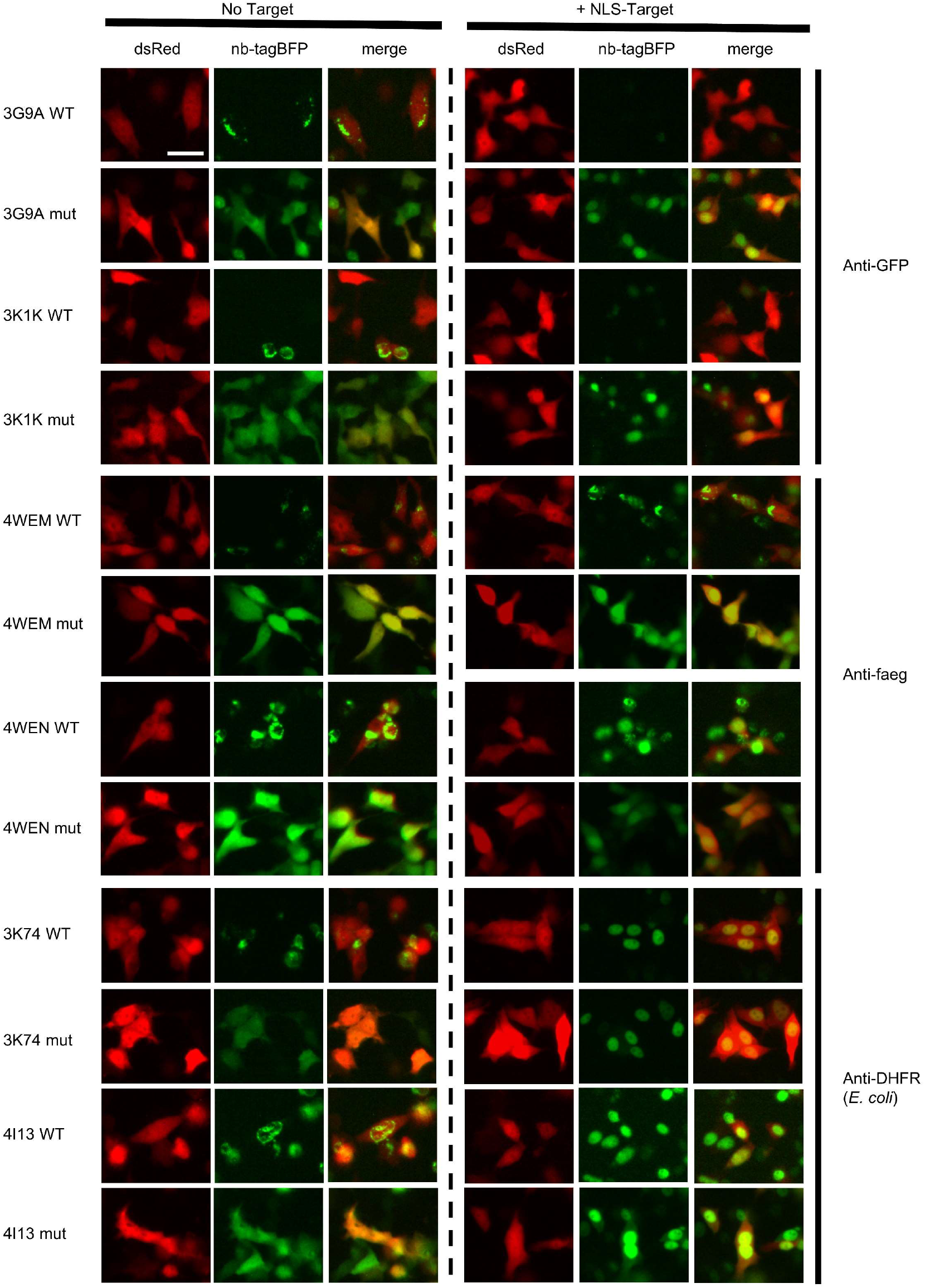
All 6 nanobodies tested for intracellular target binding. Representative images nanobody-TagBFP expression in HeLa cells in the presence and absence of nuclearly localized target for all 6 nanobodies tested. Wild-type and mutant nanobodies are depicted. Red signal is from co-transfected CAG dsRed plasmid. Transfected DNA amount was normalized by addition of empty vector plasmid to transfection mix for the “no target” condition. Scale bar is 25 μm.

